# Coevolution-based prediction of protein-protein interactions in polyketide biosynthetic assembly lines

**DOI:** 10.1101/669291

**Authors:** Yan Wang, Miguel Correa Marrero, Marnix H. Medema, Aalt D.J. van Dijk

## Abstract

Polyketide synthases are multimodular enzymes that generate diverse molecules of great pharmaceutical importance, including a range of clinically used antimicrobials and antitumor agents. Many polyketides are synthesized by type I polyketide synthases (PKSs), which are organized in assembly lines, in which multiple enzymes line up in a specific order. This order is defined by specific protein-protein interactions. The unique modular structure and catalyzing mechanism of these assembly lines makes their products predictable and also spurred combinatorial biosynthesis studies to produce novel polyketides using synthetic biology. However, predicting the interactions of PKSs, and thereby inferring the order of their assembly line, is still challenging, especially for cases in which this order is not reflected by the ordering of the PKS-encoding genes in the genome. Here, we introduce PKSpop, which uses a coevolution-based protein-protein interaction prediction algorithm to infer protein order in PKS assembly lines. Our method accurately predicts protein orders (80% accuracy). Additionally, we identify new residue pairs that are key in determining interaction specificity, and show that coevolution of N- and C-terminal docking domains of PKSs is significantly more predictive for protein-protein interactions than coevolution between ketosynthase and acyl carrier protein domains.

## Introduction

Polyketides are a structurally diverse group of specialized metabolites produced by taxonomically diverse organisms. They have a large variety of natural functions, and many of them are used in the clinic as antibiotics, chemotherapeutics, immunosuppressants or cholesterol-lowering agents ^1,2^. Many polyketides are produced by the action of modular (‘Type I’) polyketide synthases (PKSs): large multi-domain enzymes that consist of repeating modules that function in an assembly-line fashion to extend a growing polyketide chain until it is offloaded and often cyclized to form the core scaffold of the polyketide natural product ^3–5^. Each module in such an assembly line again consists of multiple domains. The three core domains are an acyltransferase (AT) domain that selects an extender unit, an acyl carrier protein (ACP) that functions as a ‘hook’ to which this unit is transferred by the AT domain, and a ketosynthase (KS) domain that catalyses the condensation reaction to extend the polyketide chain with this unit. Besides these three, PKSs may also contain ketoreductase (KR), dehydratase (DH) or enoyl reductase (ER) domains, which determine the degree to which an extender unit is reduced.

PKS proteins may consist of one or multiple modules, and a PKS assembly line frequently consists of multiple individual PKS proteins, sometimes even 5-10 in total. The final structure of the polyketide is determined by the order in which the PKS proteins interact to form the assembly line ^6^. Therefore, understanding protein-protein interactions between PKSs is of great importance. After all, accurate prediction of these interactions from sequence information is essential for predicting the chemical structure of the product, and thus makes it possible to ‘dereplicate’ the biosynthetic gene clusters (BGCs) that encode the production of polyketides to identify which of these are responsible for the production of either known or unknown (and therefore novel) molecules ^7^. Moreover, understanding and predicting these protein interactions is crucial for synthetic biology approaches to PKS engineering to be successful ^8^, as it facilitates new ways of reordering PKS assembly lines to generate novel chemistry.

Several studies have attempted to predict the order of proteins in PKS assembly lines from sequence. The simplest way of doing this is by assuming collinearity of the proteins with the order of the PKS-encoding genes in a BGC, which has frequently been observed to be true for BGCs encoding the production of known molecules ^9–11^. However, there are many cases in which the order of the PKS-encoding genes in a BGC is not compatible with collinearity (e.g., when they are located in multiple operons) ^12–16^, as shown in Figure 1. For example, in the MIBiG repository ^17^, 12 out of 133 (9%) of the multimodular type I PKS assembly lines are non-collinear. There are currently 3,302 type I PKS gene clusters in the antiSMASH database and the number increases with more and more genomes being sequenced in the future. This necessitates the use of more advanced approaches to predict protein-protein interactions. Several studies have shown that small helical domains at the N- and C-termini of PKS proteins, called docking domains, play a key role in mediating the specificity of interactions, through specific pairwise dimerization with docking domains of other proteins in the assembly line ^18–21^. Early computational analyses of the diversity of these docking domains showed that they fall into at least three compatibility classes, and that head-to-tail pairing of docking domains within the same compatibility class is necessary but not sufficient to predict the order of PKS proteins in an assembly line ^22,23^. Additionally, several studies identified potential specificity-conferring (and coevolving) residue pairs, which made it possible to predict the final order of PKS assembly lines to a larger extent ^22–24^. Still, for the large majority of assembly lines, predictions of the latest (rule-based) methods have been inconclusive, as multiple orderings are deemed equally likely ^23^.

**Figure 1.**
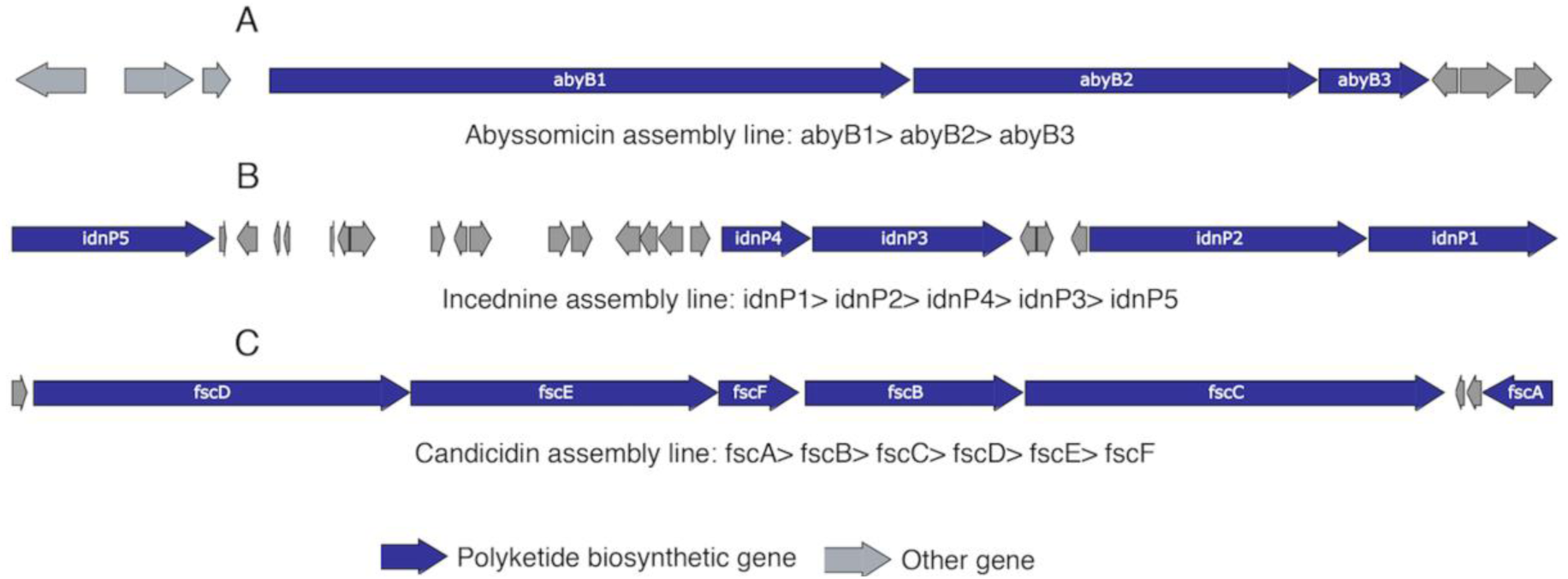
Collinear and non-collinear PKS assembly lines. In each panel, the gene arrows represent the PKS biosynthetic gene cluster; the protein order is provided underneath. (A) The protein order of the abyssomicin assembly line is collinear with the order of genes in the genome. (B) The genes in the incednine biosynthetic gene cluster span three operons. The order of the proteins in the assembly line is non-collinear with the order of the genes within the operons. (C) In the candicidin biosynthetic gene cluster the genes span more than one operon. Although there is collinearity within each operon, the order of operons is not collinear with the order of proteins in the assembly line.

Excitingly, several developments have taken place in the past years that provide new opportunities to revisit and improve predictions of PKS assembly line ordering from sequence. First, new crystal structures of cyanobacterial (‘class II’) docking domains ^25^ have revealed a markedly different tertiary structure for these domains, which shows that previous analyses based on joint alignments of multiple docking domain classes ^23^ may have been suboptimal. Second, considerable amounts of new data have been collected into standardized repositories ^17^, making it possible to substantially expand the training sets. Third, we have recently developed a new computational method that reduces noise in coevolution-based intermolecular contact predictions ^26^, leading to considerable improvements in distinguishing between protein-protein interactions and non-interactions.

Here, we provide a new prediction pipeline, PKSpop, that exploits these new insights and developments, through automated recognition of docking domain classes, coevolutionary prediction of interaction probabilities between class I docking domains using expectation-maximization with the Ouroboros method ^26^, and combining these results in a new algorithm that is usually able to predict a single most probable assembly line order. We show that PKSpop is able to accurately identify assembly line orders in non-collinear PKS systems in ∼80% of the cases. Additionally, we identify new residue pairs that are key in determining interaction specificity, and show that docking domain coevolution is significantly more predictive for protein-protein interactions than coevolution between ACP and KS domains.

## Methods and Implementation

### Dataset for learning and evaluation

Sequences of interacting docking domain pairs were obtained from The Minimum Information about a Biosynthetic Gene cluster (MIBiG) repository ^17^. 372 interacting docking domain pairs were extracted from 133 type I PKS assembly lines, whose protein orders have been established by previous studies. In addition, 15 interacting pairs were obtained from 13 assembly lines in the antiSMASH database, which stores the biosynthetic gene clusters detected and annotated by antiSMASH ^27,28^; these were all assembly lines consisting of two or three proteins, for which there was only one possible order based on the positioning of starter-AT and/or thioesterase domains at the start and end. An extra set of docking domain pairs was extracted from adjacent genes in gene clusters in the antiSMASH database. The interaction status of these pairs is unknown.

Compared to the ACP structure ^29^, the ACP domains annotated in the MIBiG and antiSMASH databases did not cover the whole ACP sequence. Thus, the sequences detected by antiSMASH as ACP domains were extended with 20 additional N-terminal residues. In addition to the interprotein KS/ACP pairs, the intra-protein pairs, whose KS and ACP domain were on the adjacent module in a protein, were extracted. The datasets were filtered by removing redundant sequences at 100% identity, as identified by CD-HIT ^30^.

### Clustering and multiple sequence alignment

Profile HMM analysis ^31^ was employed to classify and align the docking domains. It should be noted that the compatibility classes I, II and III that refer to docking domain sequences ^22^ are different from the structural classes as described by Weissman ^32^: sequence compatibility classes I, II and III correspond to structural classes 1a, 1b and 2, respectively. In our study, sequences of different compatibility classes published by Thattai et al ^22^ were aligned separately by MUSCLE ^33^ with a gap opening penalty of 11.0. Using the alignment, hmmbuild from the HMMER package ^34^ was used to build HMM profiles for each class, and hmmscan was then used to assign class membership to sequences by finding for each sequence its best match among the three HMM profiles. Subsequently, the sequences belonging to different classes were aligned against their profile by hmmalign. The conserved regions of 26 residues at the N-terminal docking domain (Ndd) and 16 residues at the end of the C-terminal docking domain (Cdd), which are involved in the protein-protein interaction ^19,21,35^, were obtained from the MSAs (Supplementary Figure 2). Docking domains used in study are described in Supplementary Dataset 1 & 2. For various alignments, the number of effective sequences (N_eff_) was calculated with conkit v0.11.2 ^36^

The type I PKS KS domains can be subdivided into those belonging to modular PKS, hybrid PKS and trans-AT PKS systems ^37–39^. As the hybrid KS domains were not included in the initial dataset, the modular KS domains were selected by matching KS domains to the modular-KS and trans-KS HMM profiles from antiSMASH ^23,40^. The ACPs that paired with the modular KS domains were then extracted as well.

### Ouroboros analysis

Datasets with different percentages of interacting pairs and different numbers of effective sequences were analyzed by Ouroboros. Since Ouroboros employs expectation-maximization (EM), each analysis was repeated three times using different random seeds to address the problem that EM might find a local optimum. In addition, different values of the int_frac parameter were used. Out of these analyses, the result with the largest log likelihood was selected. Each dataset could consist of interacting pairs, non-interacting pairs and pairs without interaction information; the assessment of performance of PPI prediction was always based on the known interacting and non-interacting pairs (note that the interaction status is not used as input by the algorithm itself).

### Selecting specificity-determining residues

Ouroboros predicts contacts by assigning each residue pair a contact score. In the structure of interacting class I docking domain pair (PDB # 1PZR), there are 31 physical contact residue pairs under the threshold of 5 ångströms ^22^. Therefore, the residue pairs of the top 31 contact scores were considered as the Ouroboros-predicted contacts and the correct prediction was defined as the intersection between the Ouroboros prediction and the physical contacts.

To find the set of residues that define the specificity of PKS PPIs, we checked the predictive performance of Ouroboros in the absence of each residue pair separately and removed the columns from the MSA corresponding to the residue pair that had the least impact on PPI prediction. With the remaining residues, we again removed the pair that had the least impact. This procedure was repeated until all residue pairs were removed. The performance was assessed by the AUC value of ROC curve, calculated by scikit-learn v0.20.2 ^41^. PyMOL v2.0 42 was used to visualize the position of selected residues in the structure^42^.

### Logistic regression model

The docking domain pairs and KS-ACP pairs were analyzed by Ouroboros separately. Then, a logistic regression model was built to predict the interaction of proteins that harbor class I docking domains. One of the predictors was the interaction probability of docking domain pairs predicted by Ouroboros; the other predictor was the interaction probability of the corresponding KS-ACP from the same protein pair. The dataset contained 80% interactions and 20% non-interactions. Five-fold cross-validation was used to test the performance of the logistic regression model.

### PKSpop: Protein order prediction pipeline

A pipeline, PKSpop was designed and implemented to predict the protein order. PKSpop takes the GenBank file of the query PKS gene cluster as detected by antiSMASH and a list of protein identifiers of the PKSs in that gene cluster as input. The code is available on http://www.bif.wur.nl/ (under ‘Software’).

PKSpop starts with extracting the docking domain sequences from the PKS proteins encoded in the cluster. First, the start protein of the assembly line is determined by locating a starter-AT domain; the end protein is determined by locating a thioesterase domain. The N- and C-terminal sequences are then assigned to one of the three different docking domain classes by hmmscan (using competitive scoring of class-specific docking domain pHMMs), and aligned using hmmalign (See Clustering and MSA section). The conserved 26-amino acid and 16-amino acid regions on the class I N- and C-termini are subsequently obtained from the alignment. Next, each N-terminal sequence is paired with all C-terminal sequences. Two paired fasta files, one of N-terminal sequences and one of C-terminal sequences, are generated by integrating the query sequences with 222 interacting sequences and 301 sequences from adjacent genes in gene clusters. The two fasta files with paired docking domain sequences are then input into Ouroboros and analysed under user-defined int_frac parameters (default [0.80, 0.90]) and a user-indicated number of times the analysis is repeated.

Ouroboros then outputs the pairwise interaction probabilities of the query docking domain pairs from the result with the largest log-likelihood. These are filled into an interaction probability matrix, where the C-termini are in rows and the N termini are in columns. From the matrix, the pairs that do not belong to class I are first matched to each other. (If there are multiple C- and N-termini of class II or III, multiple possible assembly line orders will be elaborated in parallel.) Then, the class I pair with the highest probability is found and fixed if it is not conflict with the already fixed pairs. If there is more than one pair that has the same highest probability, multiple possibilities will be accounted for again. The steps are repeated until all the values in the matrix have been traversed, or the procedure stops earlier if an order that consists of all the query proteins is predicted. If no assembly line order results that covers all query PKS proteins, the process finishes without a result (Supplementary Figure 1).

## Result & Discussion

### A new method to predict PKS order in assembly line

Predicting protein order in polyketide assembly lines is different from the typical pairwise PPI studies. This is because of the nature of the assembly line, in which a protein can only interact with one upstream and one downstream protein. To infer protein order in assembly lines, PKSpop comprises three main steps: 1) Identify class memberships for query docking domains and align the sequences; 2) Pair each class I N-terminal docking domain (Ndd) with all class I C-terminal docking domains (Cdds) and use coevolution to predict the interaction probability for all possible pairs and enter these into a probability matrix; 3) Infer the protein order by a greedy probability matrix-filling method which takes the assembly line constraints and compatibility class into account.

We first set out to automatically classify docking domains into classes, and to assess for which of these classes sufficient data is available to predict their protein-protein interactions (PPIs). The performance of coevolution-analysis-based PPI prediction depends highly on the alignment quality and shifts in interaction site can lead to mispredictions ^26,43^. While a previous method ^23^ used joint alignment of the docking domains from all three compatibility classes ^22^, new structural studies have shown that these docking domain classes have a markedly different tertiary structure and therefore should not be aligned ^19,21,25^. Instead, it is better to analyze the classes separately. To annotate docking domains by class, we built HMM profiles of all three docking domain compatibility classes from sequences published by Thattai et al ^22^. Then, 384 interacting Cdd and Ndd pairs were clustered by finding for each domain its best match among the three HMM profiles. As a result, 222 class I and 85 class II interacting docking domain pairs were obtained. There were 64 pairs that contained at least one class III docking domain; 27 of these involved cases where a single class III docking domain was paired with a domain which was not assigned to any class. One possible reason is that the number of sequences used to build class III HMM profiles was only 14, which might be too little to cover the sequence diversity of class III docking domains. The other possible reason might be that class III was not correctly defined: in previous clustering ^22^, most pairs that were ostensibly mismatched based on comprising members of different classes have one of their sequences assigned into class III, and, in the sequence similarity networks, class III appears to be much less coherent then classes I and II. Also, the docking domains clustered in sequence class III according to previous studies ^22,43^ (which corresponds to structural class 2 ^25^) have dissimilar structures, as can be seen from the structures of CurK/CurL (PDB # 4MYY) and CurG/CurH (PDB # 4MYZ) ^25^. Therefore, it is still an open question whether there is a sufficiently homogeneous “class III” that truly represents a group of structurally similar docking domains across organisms.

Using the alignments of the class I docking domains, we applied Ouroboros, our recently developed method to predict protein interactions based on intermolecular coevolution.. Ouroboros takes only the alignments as input and generates pairwise protein interaction probabilities of the class I docking domains. Testing various input datasets, we demonstrate that coevolutionary analysis can be successfully applied to predict class I docking domain interactions. Before presenting these results, we first give an overall overview of PKSpop, the pipeline we developed to apply coevolutionary analysis to PKS assembly lines.

### PKSpop predicts PKS protein orders with high accuracy

To assess the performance of the PKSpop pipeline specifically in the context of assembly lines that do not follow the collinearity rule, the following ten assembly lines were investigated: BE-14106, candicidin, FK-520, incednine, kendomycin, kijanimicin, lobophorin, ML-449, nanchangmycin, and oligomycin. In these ten assembly lines, there are 44 interacting protein pairs among 262 pair combinations. Of these, the order of 8 lines (80% accuracy; nanchangmycin and oligomycin were the exceptions) and the interactions of 36 protein pairs (82% accuracy) were correctly predicted. Figure 2 shows an example of predicting the protein order of the candicidin assembly line. FscA is the first protein as it has an initiating AMP-binding domain, and fscF is the final protein as it does not have a Cdd (Figure 2A). The Cdd of fscD (fscD-Cdd) and Ndd of fscE (fscE-Ndd) were clustered into class II, while other docking domains all belong to class I. Hence, fscD and fscE were first connected, as fscD-Cdd and fscE-Ndd comprised the only pair that does not belong to class I. The pairwise interaction probabilities of the docking domains were predicted by Ouroboros and filled into the matrix (Figure 2B). Based on the probability matrix, fscF was then connected to fscE, since fscE-Cdd and fscF-Ndd have the highest interaction probability in the matrix. Subsequently, the interaction of other Cdds with fscF-Ndd was ruled out. Iteratively, the pair with the highest probability in the remaining rows and columns was identified and connected repeatedly to predict the final (correct) order (Figure 2C).

**Figure 2.**
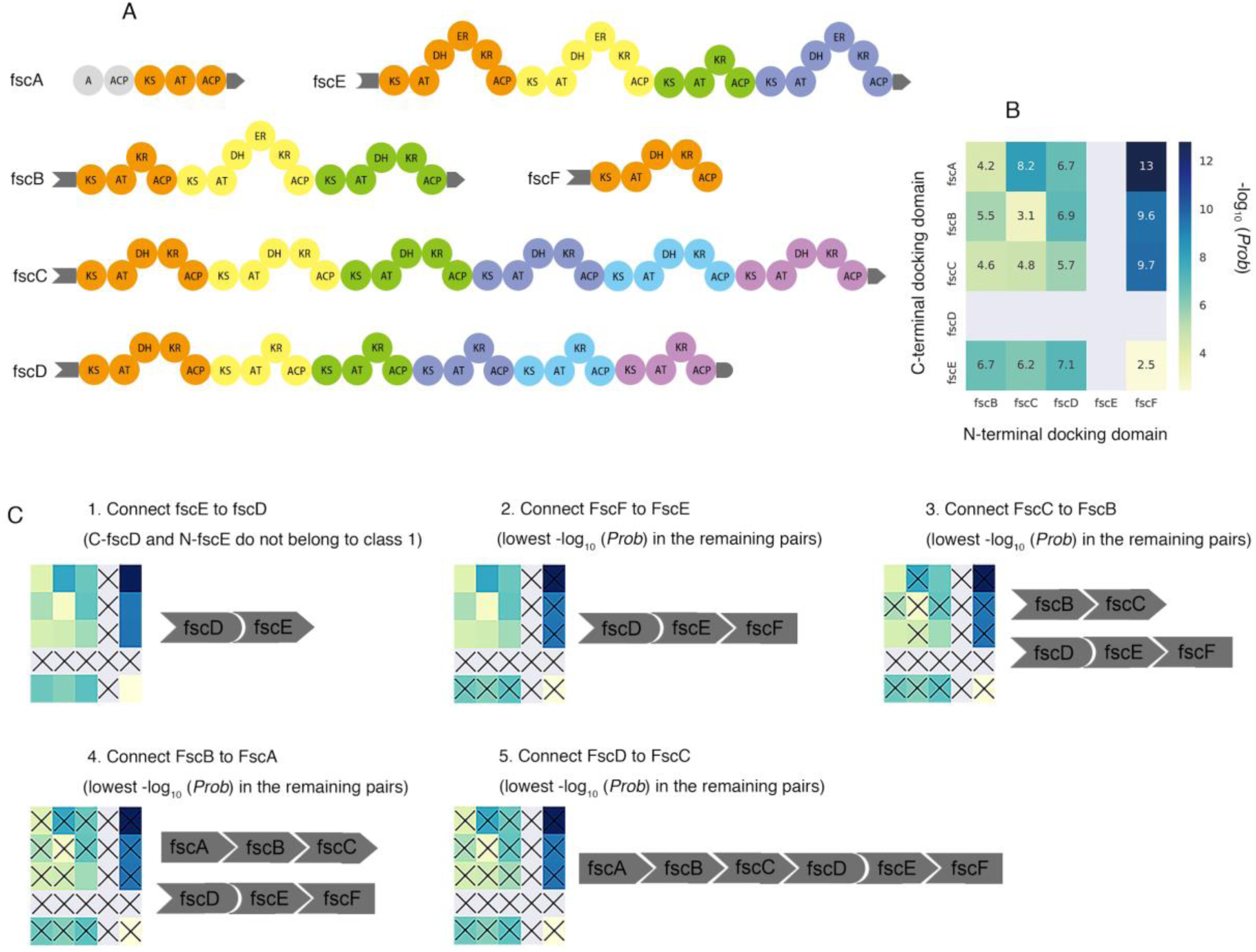
Predicting the protein order of Candicidin assembly line. (A) The PKS proteins and their domains in the Candicidin assembly line. (B) Interaction of PKSs predicted by Ouroboros. Each cell represents a possible interaction of a docking domain pair and is coloured by the value of −log_10_(Interaction probability). Cells of C-fscD and N-fscE are colored grey, as these docking domains do not belong to class I. (C) PKSpop procedure to predict the protein order from the interaction probabilities. First, fcsD and fscE are connected, as their docking domains do not belong to class I. Subsequently, pairs of docking domains are iteratively connected by finding the matrix cell with the highest probability, while crossing out interactions that are no longer possible. In the end, a single predicted order remains.

We compared the performance of our algorithm to a previously published method ^23^, which uses two residue pairs as a “docking code” to infer protein orders. With this method, five assembly lines out of ten were correctly predicted. Moreover, it should be noted that our method outputs a single correct order for all eight assembly lines. In contrast, the previous method predicted multiple possible orders with the same highest score for each assembly line (20 for BE-14106, 30 for candicidin, 6 for kijanimicin, 4 for lobophorin and 20 for ML-449), which reiterates the advantage of our probabilistic approach.

### Identification of specificity-conferring residue pairs in class I docking domains

Based on structural information, there are 31 contacting residue pairs on class I docking domains. We selected the top 31 residue pairs scored by Ouroboros as involved in intermolecular coevolution, and of these, 21 were involved in intermolecular contacts (p-value 7.47e-15, Fisher exact test). To study the impact on PPI specificity of the selected residue pairs, MSAs were generated in which all the predicted residues were removed. The prediction of PPIs based on the remaining residues failed (AUC=0.55), which indicates that the determinant residue pairs lie in the selected pairs. By removing the least important contact residue pairs one by one from the MSA, we found that performance decreased drastically after pairs C11-N9, C12-N5 and C4-N3 were removed (Supplementary Figure 3). Removing all three pairs still led to a model with more predictive power (AUC=0.67) than a model in which all contacting pairs were removed. However, when using the three pairs alone, the predictive performance (AUC=0.83) is similar to that of using the whole sequence (AUC=0.81) (Supplementary Table 1). This indicates that they are likely to be specificity-determining residues (Figure 3).

**Figure 3.**
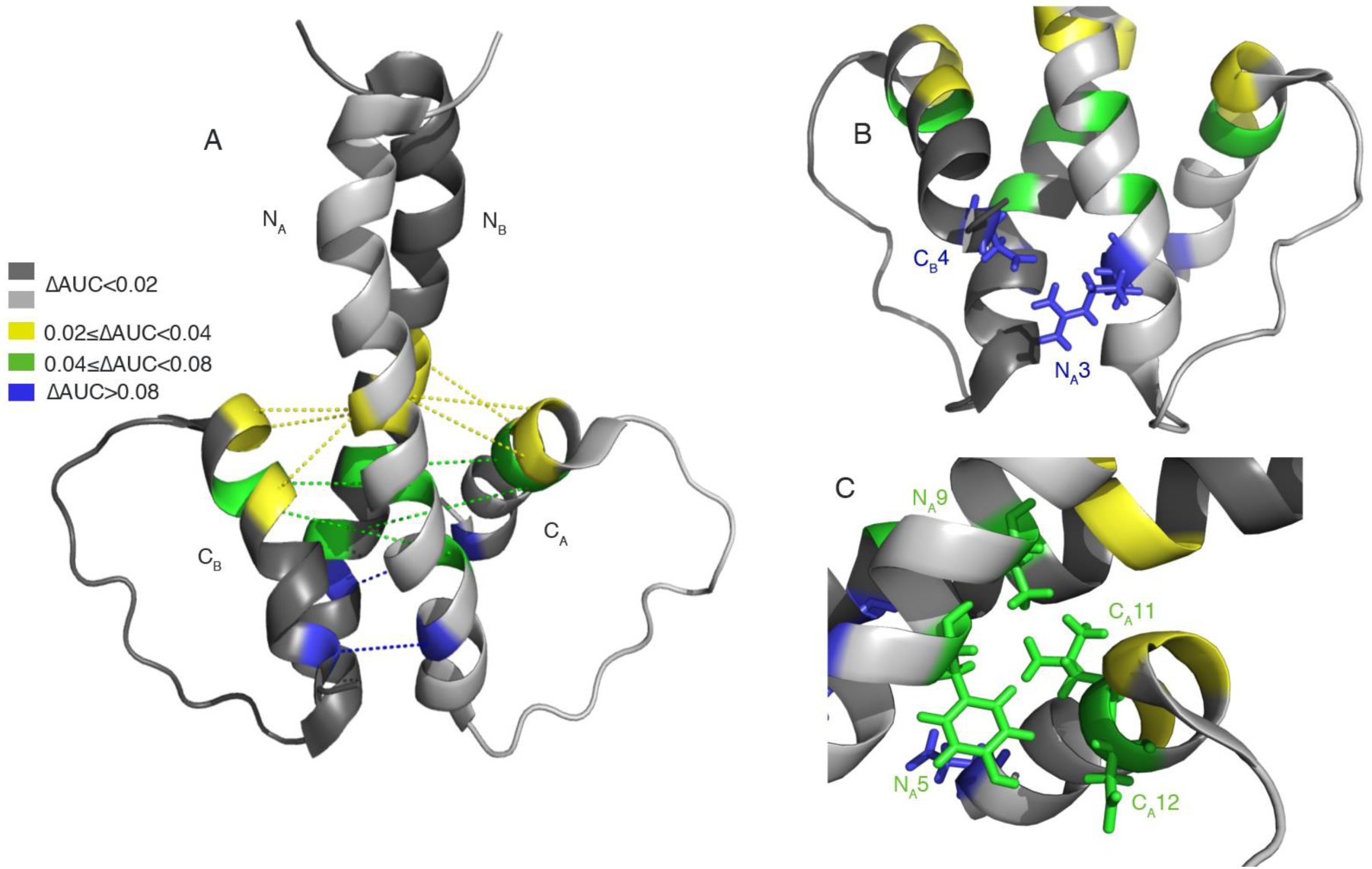
Docking domain interaction structure with predicted crucial residues. (A) The docking domain interaction structure (PDB # 1PZR). PKS proteins are homodimers and therefore there are two docking domain pairs, C_A_-N_A_ and C_B_-N_B_. Residue pairs (linked by dashed lines) are coloured according to the decrease of AUC after they were removed from the sequences. (B) The physical position of residue pair C4-N3, which was predicted to contribute to protein interaction specificity by Ouroboros. (C) The physical position of C12-N5 and C11-N9, which were also predicted to contribute to protein interaction specificity. The C12-N5 pair was found for the first time in our study as a determinant of PPI specificity.

Next, we analyzed how the three specificity-determining residue pairs influence the order prediction (Supplementary Table 1). When using the three pairs to infer protein order, the correct prediction was 8/10, which was the same as that of using the whole sequence. Although using only one of the pairs can still lead to a similar accuracy, in this case not always a single unique order was predicted. Among the three pairs, C4-N3 showed the greatest importance in PPI specificity, as using it alone could achieve a highest AUC on PPI prediction and a highest accuracy on order prediction. While C4-N3 and C11-N9 were also identified as the protein-contacting parts of the “code word” of PPI specificity by Thattai et al ^22^, the C12-N5 pair is found for the first time in our study as a determinant of PPI specificity.

### Docking domains instead of KS-ACP domain pairs contribute to predicting PPI

To investigate whether KS-ACP domain pairs can also contribute to predicting PPI specificity in PKSs, a logistic regression model was used to predict the interaction of PKS proteins that harbor a class I docking domain. One of the predictors was the interaction probability of docking domain pairs predicted by Ouroboros. The other predictor was the interaction probability of the corresponding KS-ACP pairs, which have also been shown to play a role in mediating protein-protein interactions between PKSs ^8,44,45^. Different training and testing sets were created by 5-fold cross-validation, and the ROC curve was plotted on each testing set (Supplementary Figure 4). The resulting AUC of 0.83±0.02 demonstrates that the model is predictive of PKS protein interactions. In the logistic model, the coefficient of the KS-ACP interaction probability was around −0.27±0.16, much lower than that of docking domain, 2.98±0.09, which indicates that the Ouroboros’ result on KS-ACP does not contribute to the PPI prediction. To further evaluate this result, interaction probabilities of the two domain pairs obtained from Ouroboros were used separately to predict the PPI (Figure 4). The ROC curves of docking domains have similar shape and AUCs with that of logistic regression model, while that of KS-ACP are not better than random guess.

**Figure 4.**
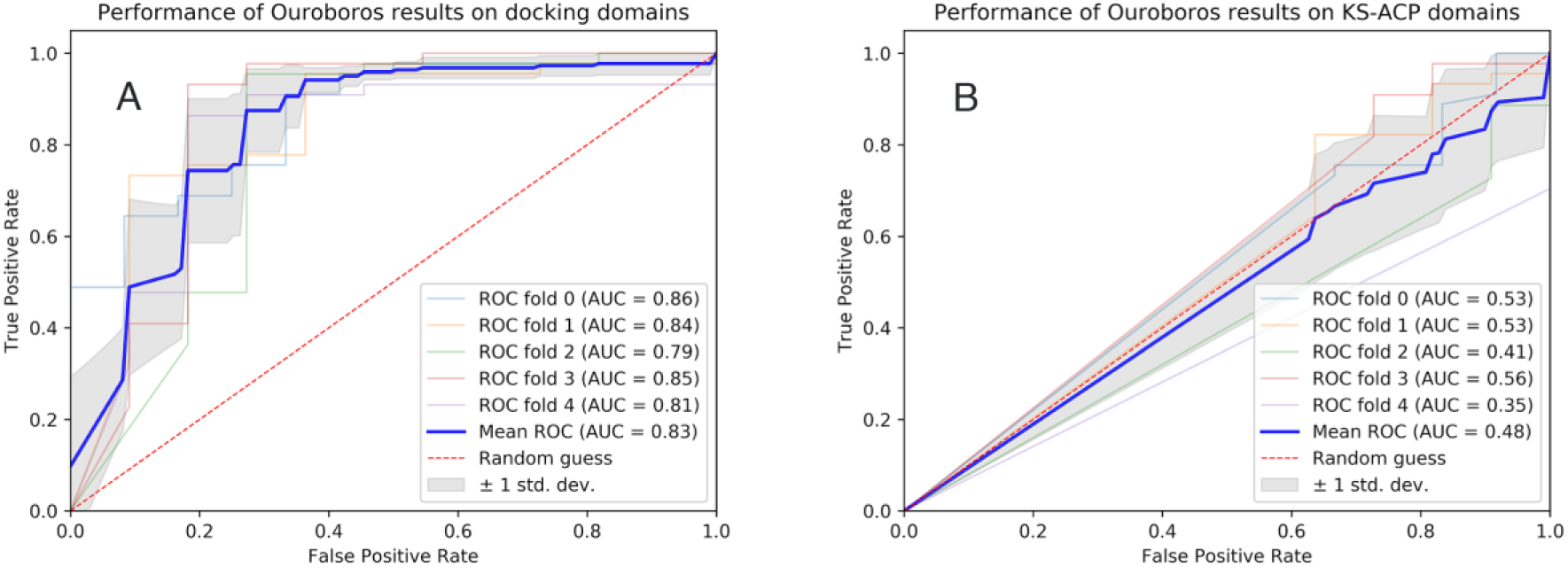
PKS protein interaction prediction performance. Receiver operating characteristic (ROC) curves of logistic models based on inferred interaction probabilities, using only (A) class I docking domains and (B) KS-ACP domain sequences to predict the PPIs.

The poor predictive ability of Ouroboros result on KS-ACP pairs might indicate that coevolution is not suitable to predict KS-ACP interaction. According to previous studies ^45,46^, R23 on DEBS2 ACP helix I was found to be the determinant of inter-protein KS-ACP interaction specificity by mutagenesis ^45^, but the contact score of this residue to any other KS residues is relatively low in the Ouroboros result. The low contact score might mean that contact residues in KS domains are evolutionary conserved and therefore cannot be detected by a coevolution-based method. The poor predictive ability might also indicate that KS-ACP interaction is not involved in mediating the specificity of PPIs between PKSs, in contrast to previous assumptions ^8^. Circumstantial evidence for this is given by the DEBS assembly line, for which functional protein interaction only requires compatible docking domain pairs ^6,47^.

### Noise level and sequence variation influence the prediction

Predicting the protein order in assembly lines is based on predictions of protein-protein interactions by Ouroboros, and this prediction can be influenced by the noise level (number of non-interacting protein pairs) and sequence variation of the input proteins ^26^. To investigate the sensitivity of our method, we therefore tested Ouroboros with different docking domain datasets.

To compare the predictive performance with different input noise levels (number of non-interacting protein pairs), five datasets with 80% interacting class I docking domains and three datasets with 60% interacting domains were generated. All datasets comprised the same 222 interacting domain pairs and a number of different non-interacting domain pairs. Sequence variation was measured by the number of effective sequences (N_eff_, number of sequences in a dataset whose pairwise identities are below 90%). To increase N_eff_, extra docking domain pairs were added to the datasets of 80% and 60% interacting pairs. The 301 extra pairs were generated by pairing the docking domains from adjacent genes in polyketide biosynthetic gene clusters compatible with collinearity. According to the collinearity rule, these extra data are likely to include a high percentage of interacting pairs and can be used to increase the sequence variation without introducing too much noise. The interaction information of these extra pairs is unknown, and the performance assessment was based on the known interacting and non-interacting pairs. Importantly, the knowledge on interaction status was not used for training the method, only for evaluating its performance. Table 1 shows Ouroboros’ predictive performance, measured by the Matthew correlation coefficient, on the different datasets. The results clearly demonstrate that with more interacting domain pairs and greater N_eff_ in the dataset, Ouroboros tends to perform better. Therefore, when using Ouroboros to generate the pairwise interaction matrix in the order prediction pipeline, the query sequences are combined with all the existing interacting protein pairs and the extra pairs.

**Table 1.** Predictive performance on datasets with different percent of interacting pairs and different number of effective sequences. The N_eff_ and Matthew correlation coefficient (MCC) are the mean values of 3 (with 60% interacting pairs) or 5 (with 80% interacting pairs) datasets.

| Dataset composition: % of interacting pairs (+ extra pairs) | $N_{\text{eff}}$ | MCC |
| --- | --- | --- |
| 60% | 212±1 | 0.22±0.02 |
| 60% + extra pairs | 468±4 | 0.45±0.06 |
| 80% | 211±3 | 0.30±0.05 |
| 80% + extra pairs | 465±2 | 0.60±0.07 |

Since Ouroboros is able to infer interaction of proteins that contain class I docking domains with good accuracy, it was then applied to class II docking domains. Lacking equal amounts of sequence data, an MSA with a N_eff_ of only 168 was created, which comprised a set of 80% interacting pairs and a set of extra pairs without interaction information. The result shows that Ouroboros failed to perform well on the class II docking domains with this amount of effective sequences (Supplementary Figure 5). To analyze whether the poor performance is caused by the lack of sequence variation, datasets of class I docking domain with similar N_eff_ were created and input into Ouroboros. The performance on class I domains is indeed better, but the standard deviation is relatively high compared to that obtained with the higher N_eff_. Altogether, this suggests that the performance is unstable on such a small dataset. With more PKS biosynthetic gene clusters being sequenced in the future, there will be more class II docking domains available, which can increase the sequence variation in MSA and may help improve the prediction for this class.

The fact that currently Ouroboros cannot accurately predict protein interactions for class II or class III docking domains, means that PKSpop is currently unable to infer assembly lines whose docking domains all belong to class II or III. However, as demonstrated above, the PKSpop pipeline can work on assembly lines that contain limited numbers of class II or III docking domain pairs in addition to class I docking domain pairs, by connecting the docking domains of class II and/or III, respectively. Based on the composition of docking domain classes in BGCs currently deposited in MIBiG, we estimate that PKSpop is able to predict assembly line orders in ∼90% of the cases.

### Conclusions and future perspectives

We have presented a method to predict protein order in PKS biosynthetic assembly lines. Our method, PKSpop, enables to deal with the fact that often, gene order in a gene cluster is not predictive of the protein interaction order in the assembly line. The basis of our approach is our recently developed coevolution-based method for protein interaction prediction, which assigns a likelihood of interaction to pairs of proteins based on whether coevolution is predicted between these proteins. It does so without requiring any knowledge on protein interactions or residue contacts. Here, we presented a greedy matrix-filling approach to convert the coevolution based interaction probability between multiple proteins to an ordering of proteins in an assembly line. Application of this approach to PKS gene clusters demonstrated its performance. This now enables to predict protein-protein interactions and hence putative core scaffold chemical structures for clusters which defy the collinearity rule; hitherto, interactions in these clusters could not well be predicted. In the near future, we plan to incorporate PKSpop as a prediction feature into the antiSMASH pipeline ^40^.

In addition, our method identified specific residue pairs, one of which was highlighted for the first time as specificity-determining; this pair might be further studied to provide insight into the determinants of PKS specificity. Finally, in contrast to collinearity-based predictions, our approach enables predicting interactions between proteins from different assembly lines. This will be of great use for efforts to engineer synthetic PKS assembly lines ^32,48,49^ consisting of entirely new combinations of proteins.

## Supplements

**Dataset 1.** Source of docking domains from MIBiG database, including MIBiG BGC identifier, protein identifier, polyketide biosynthetic assembly line and organism.

**Dataset 2.** Source of docking domains from antiSMASH database, including GenBank accession, antiSMASH gene cluster number and organism.

**SI Figure 1.**
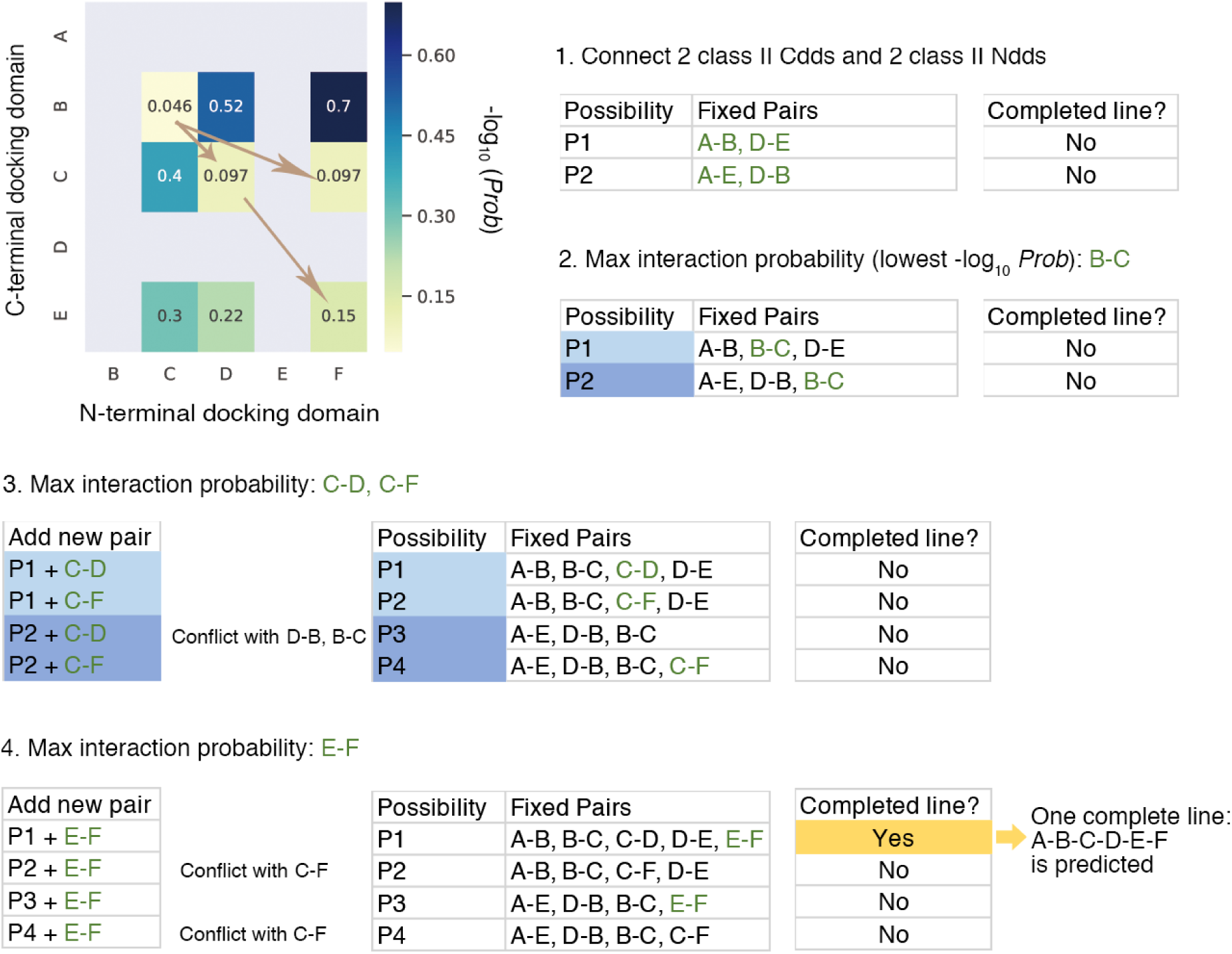
Example of predicting protein order from interaction probability matrix. The protein pairs, for example the A-B, refers to the interaction of the C-terminal docking domain of protein A with the N-terminal docking domain of protein B.

**SI Figure 2.**
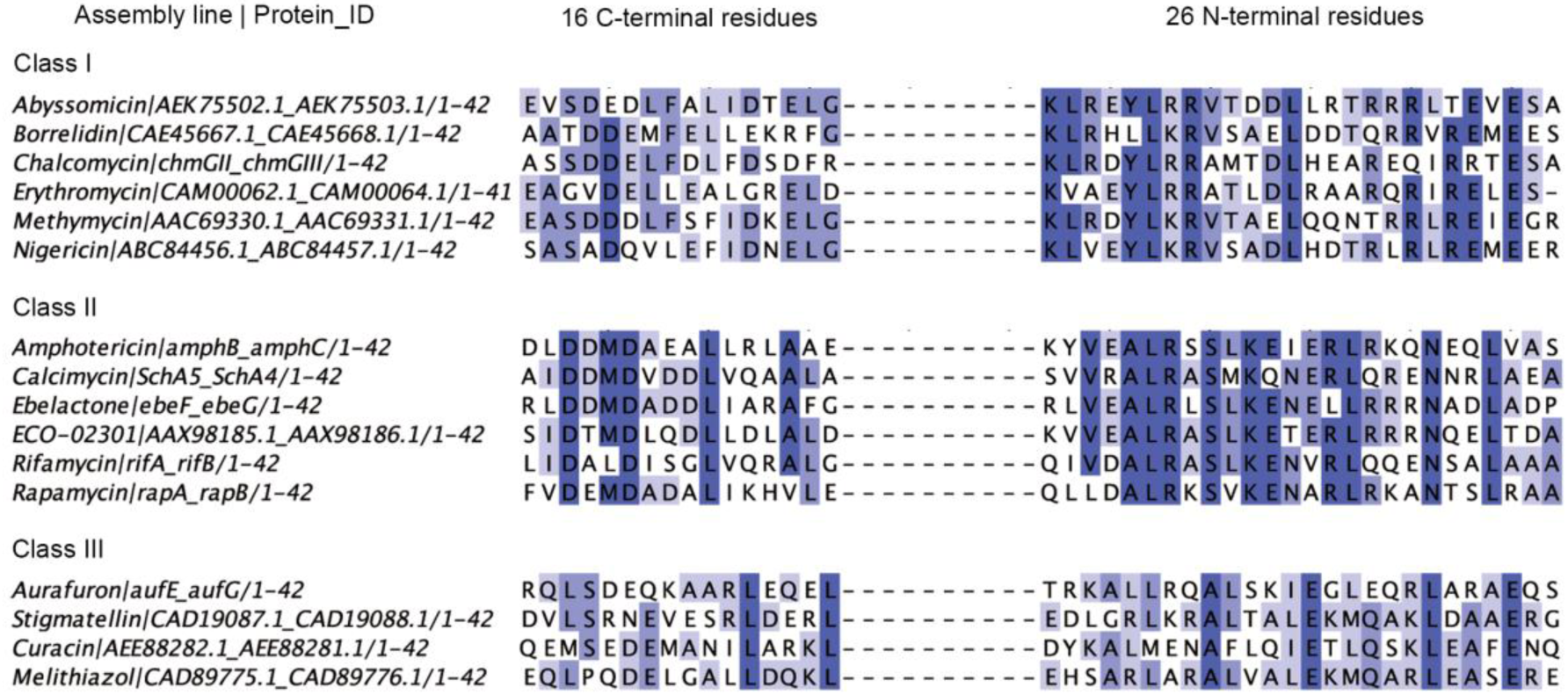
Example of multiple sequence alignment of conserved regions on docking domains of 3 compatibility classes. Each line represents an interacting docking domain pair. The labels consist of the polyketide biosynthetic assembly lines and the IDs of proteins which harbor the docking domains. Residues are coloured by identity in MSA (darker colour for higher identity).

**SI Figure 3.**
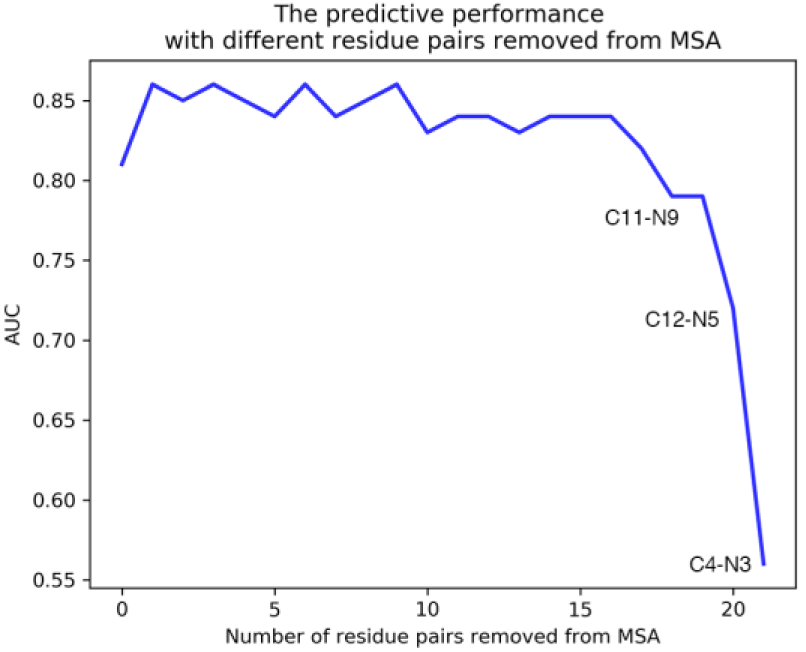
Predictive performance of removing the least important residue pairs one by one from the MSA. Residue pairs which caused drastic decrease of AUC score are indicated.

**SI Figure 4.**
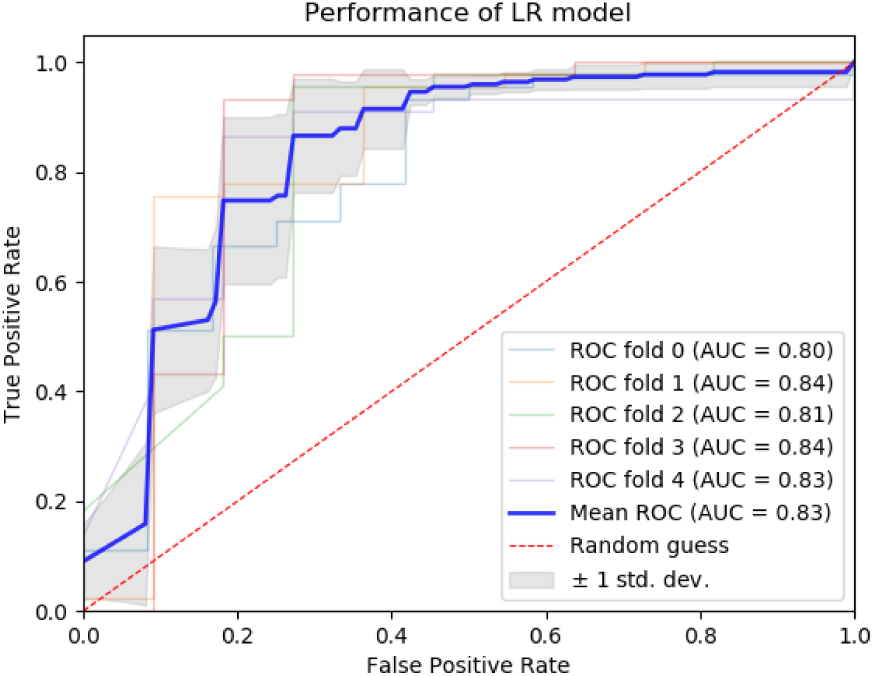
Performance of logistic regression model built on Ouroboros results of class 1 docking domain pairs and the corresponding KS-ACP pairs. The light curves show performances of model on 5 testing sets; the bold blue curve presents the averaged performance; the grey area is the estimated ROC ±1 standard deviation.

**SI Figure 5.**
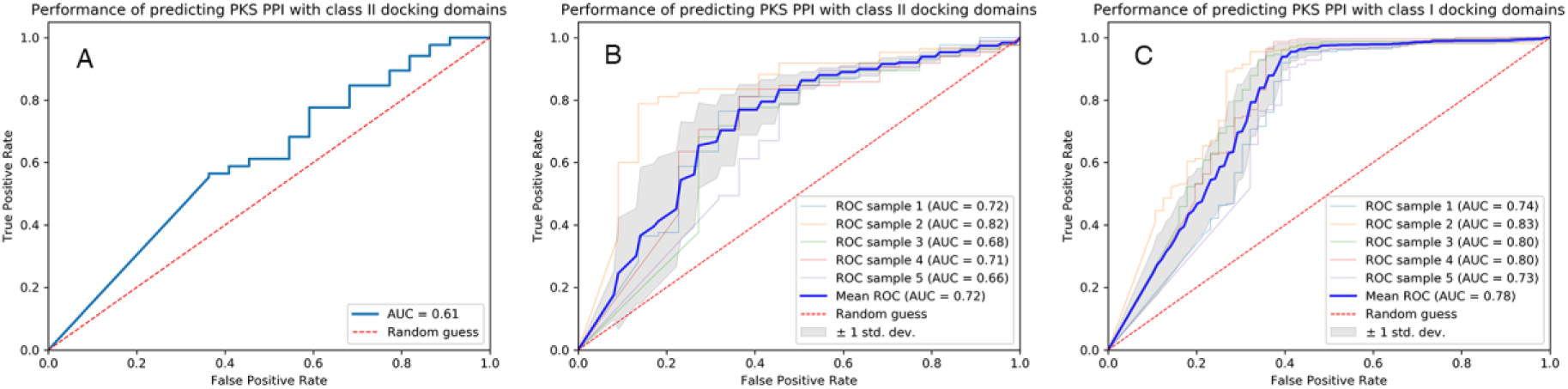
Predictive performance on (A) one dataset of class II docking domains (N_eff_=168), (B) five datasets on class I docking domains (N_eff_=168±1) and C) five datasets of class I docking domains (N_eff_=465±2).

**SI Table 1.** Performance of using different residue pairs to predict the PPIs and protein order in polyketide assembly lines. The first column is the residue pairs used. The second column is the AUC score of the PPI prediction. The third column is the accuracy of protein order prediction using the corresponding pairs and the last column is the number of unique predictions among all correctly predicted orders.

| Residue pair used to predict interaction | AUC of predicting protein interaction | Correct rate of predicting protein order | Unique order predicted |
| --- | --- | --- | --- |
| All | 0.81 | 8/10 | 8/8 |
| C4-N3, C12-N5, C11-N9 | 0.83 | 8/10 | 8/8 |
| C4-N3 | 0.80 | 8/10 | 7/8 |
| C11-N9 | 0.66 | 7/10 | 4/7 |
| C12-N5 | 0.61 | 8/10 | 4/8 |

## References

1. Demain, A. L. Importance of microbial natural products and the need to revitalize their discovery. J. Ind. Microbiol. Biotechnol. 41, 185–201 (2014).

2. Newman, D. J. & Cragg, G. M. Natural Products as Sources of New Drugs from 1981 to 2014. J. Nat. Prod. 79, 629–661 (2016).

3. Fischbach, M. A. & Walsh, C. T. Assembly-line enzymology for polyketide and nonribosomal Peptide antibiotics: logic, machinery, and mechanisms. Chem. Rev. 106, 3468–3496 (2006).

4. Dutta, S. et al. Structure of a modular polyketide synthase. Nature 510, 512–517 (2014).

5. Robbins, T., Liu, Y.-C., Cane, D. E. & Khosla, C. Structure and mechanism of assembly line polyketide synthases. Curr. Opin. Struct. Biol. 41, 10–18 (2016).

6. Dodge, G. J., Maloney, F. P. & Smith, J. L. Protein–protein interactions in ‘cis-AT’polyketide synthases. Natural Product Reports 35, 1082–1096 (2018).

7. Dejong, C. A. et al. Polyketide and nonribosomal peptide retro-biosynthesis and global gene cluster matching. Nat. Chem. Biol. 12, 1007–1014 (2016).

8. Klaus, M. et al. Protein-Protein Interactions, Not Substrate Recognition, Dominate the Turnover of Chimeric Assembly Line Polyketide Synthases. J. Biol. Chem. 291, 16404–16415 (2016).

9. Donadio, S., Staver, M. J., McAlpine, J. B., Swanson, S. J. & Katz, L. Modular organization of genes required for complex polyketide biosynthesis. Science 252, 675–679 (1991).

10. Donadio, S. & Katz, L. Organization of the enzymatic domains in the multifunctional polyketide synthase involved in erythromycin formation in Saccharopolyspora erythraea. Gene 111, 51–60 (1992).

11. Yu, T.-W. et al. Direct evidence that the rifamycin polyketide synthase assembles polyketide chains processively. Proceedings of the National Academy of Sciences 96, 9051–9056 (1999).

12. Jørgensen, H. et al. Biosynthesis of macrolactam BE-14106 involves two distinct PKS systems and amino acid processing enzymes for generation of the aminoacyl starter unit. Chem. Biol. 16, 1109–1121 (2009).

13. Takaishi, M., Kudo, F. & Eguchi, T. Identification of the incednine biosynthetic gene cluster: characterization of novel β-glutamate-β-decarboxylase IdnL3. J. Antibiot. 66, 691–699 (2013).

14. Zhang, H. et al. Elucidation of the kijanimicin gene cluster: insights into the biosynthesis of spirotetronate antibiotics and nitrosugars. J. Am. Chem. Soc. 129 14670–14683 (2007).

15. Sun, Y. et al. A complete gene cluster from Streptomyces nanchangensis NS3226 encoding biosynthesis of the polyether ionophore nanchangmycin. Chem. Biol. 10, 431–441 (2003).

16. Wenzel, S. C., Bode, H. B., Kochems, I. & Müller, R. A type I/type III polyketide synthase hybrid biosynthetic pathway for the structurally unique ansa compound kendomycin. Chembiochem 9, 2711–2721 (2008).

17. Medema, M. H. et al. Minimum Information about a Biosynthetic Gene cluster. Nat. Chem. Biol. 11, 625–631 (2015).

18. Tsuji, S. Y., Cane, D. E. & Khosla, C. Selective protein-protein interactions direct channeling of intermediates between polyketide synthase modules. Biochemistry 40, 2326–2331 (2001).

19. Broadhurst, R. W., Nietlispach, D., Wheatcroft, M. P., Leadlay, P. F. & Weissman, K. J. The structure of docking domains in modular polyketide synthases. Chem. Biol. 10, 723–731 (2003).

20. Weissman, K. J. The structural basis for docking in modular polyketide biosynthesis. Chembiochem 7, 485–494 (2006).

21. Buchholz, T. J. et al. Structural basis for binding specificity between subclasses of modular polyketide synthase docking domains. ACS Chem. Biol. 4, 41–52 (2009).

22. Thattai, M., Burak, Y. & Shraiman, B. I. The origins of specificity in polyketide synthase protein interactions. PLoS Comput. Biol. 3, 1827–1835 (2007).

23. Yadav, G., Gokhale, R. S. & Mohanty, D. Towards Prediction of Metabolic Products of Polyketide Synthases: An In Silico Analysis. PLoS Computational Biology 5, e1000351 (2009).

24. Burger, L. & van Nimwegen, E. Accurate prediction of protein-protein interactions from sequence alignments using a Bayesian method. Mol. Syst. Biol. 4, 165 (2008).

25. Whicher, J. R. et al. Cyanobacterial polyketide synthase docking domains: a tool for engineering natural product biosynthesis. Chem. Biol. 20, 1340–1351 (2013).

26. Correa Marrero, M., Immink, R. G. H., de Ridder, D. & van Dijk, A. D. J. Improved inference of intermolecular contacts through protein–protein interaction prediction using coevolutionary analysis. Bioinformatics (2019). doi:10.1093/bioinformatics/bty924

27. Blin, K., Medema, M. H., Kottmann, R., Lee, S. Y. & Weber, T. The antiSMASH database, a comprehensive database of microbial secondary metabolite biosynthetic gene clusters. Nucleic Acids Res. 45, D555–D559 (2017).

28. Weber, T. et al. antiSMASH 3.0—a comprehensive resource for the genome mining of biosynthetic gene clusters. Nucleic Acids Research 43, W237–W243 (2015).

29. Alekseyev, V. Y., Liu, C. W., Cane, D. E., Puglisi, J. D. & Khosla, C. Solution structure and proposed domain-domain recognition interface of an acyl carrier protein domain from a modular polyketide synthase. Protein Science 16, 2093–2107 (2007).

30. Li, W. & Godzik, A. Cd-hit: a fast program for clustering and comparing large sets of protein or nucleotide sequences. Bioinformatics 22, 1658–1659 (2006).

31. Eddy, S. R. Profile hidden Markov models. Bioinformatics 14, 755–763 (1998).

32. Weissman, K. J. Genetic engineering of modular PKSs: from combinatorial biosynthesis to synthetic biology. Nat. Prod. Rep. 33, 203–230 (2016).

33. Edgar, R. C. MUSCLE: multiple sequence alignment with high accuracy and high throughput. Nucleic Acids Res. 32, 1792–1797 (2004).

34. Eddy, S. R. Accelerated Profile HMM Searches. PLoS Comput. Biol. 7, e1002195 (2011).

35. Weissman, K. J. Single amino acid substitutions alter the efficiency of docking in modular polyketide biosynthesis. Chembiochem 7, 1334–1342 (2006).

36. Simkovic, F., Thomas, J. M. H. & Rigden, D. J. ConKit: a python interface to contact predictions. Bioinformatics 33, 2209–2211 (2017).

37. Miyanaga, A., Kudo, F. & Eguchi, T. Protein-protein interactions in polyketide synthase-nonribosomal peptide synthetase hybrid assembly lines. Nat. Prod. Rep. 35, 1185–1209 (2018).

38. Scotti, C. et al. A Bacillus subtilis large ORF coding for a polypeptide highly similar to polyketide synthases. Gene 130, 65–71 (1993).

39. Piel, J. Biosynthesis of polyketides by trans-AT polyketide synthases. Nat. Prod. Rep. 27, 996–1047 (2010).

40. Blin, K. et al. antiSMASH 4.0-improvements in chemistry prediction and gene cluster boundary identification. Nucleic Acids Res. 45, W36–W41 (2017).

41. Varoquaux, G. et al. Scikit-learn. GetMobile: Mobile Computing and Communications 19, 29–33 (2015).

42. Support pymol.org. Available at: https://pymol.org/2/support.html?#citing. (Accessed: 16th April 2019)

43. Uguzzoni, G. et al. Large-scale identification of coevolution signals across homooligomeric protein interfaces by direct coupling analysis. Proceedings of the National Academy of Sciences 114, E2662–E2671 (2017).

44. Weissman, K. J., Hong, H., Popovic, B. & Meersman, F. Evidence for a protein-protein interaction motif on an acyl carrier protein domain from a modular polyketide synthase. Chem. Biol. 13, 625–636 (2006).

45. Kapur, S., Chen, A. Y., Cane, D. E. & Khosla, C. Molecular recognition between ketosynthase and acyl carrier protein domains of the 6-deoxyerythronolide B synthase. Proc. Natl. Acad. Sci. U. S. A. 107, 22066–22071 (2010).

46. Kapur, S. et al. Reprogramming a module of the 6-deoxyerythronolide B synthase for iterative chain elongation. Proc. Natl. Acad. Sci. U. S. A. 109, 4110–4115 (2012).

47. Wu, N., Cane, D. E. & Khosla, C. Quantitative analysis of the relative contributions of donor acyl carrier proteins, acceptor ketosynthases, and linker regions to intermodular transfer of intermediates in hybrid polyketide synthases. Biochemistry 41, 5056–5066 (2002).

48. Poust, S., Hagen, A., Katz, L. & Keasling, J. D. Narrowing the gap between the promise and reality of polyketide synthases as a synthetic biology platform. Curr. Opin. Biotechnol. 30, 32–39 (2014).

49. Hertweck, C. Decoding and reprogramming complex polyketide assembly lines: prospects for synthetic biology. Trends Biochem. Sci. 40, 189–199 (2015).

